# A chemiluminescent protease probe for rapid, sensitive, and inexpensive detection of live *Mycobacterium tuberculosis*

**DOI:** 10.1101/2020.09.14.296772

**Authors:** Brett M. Babin, Gabriela Fernandez-Cuervo, Jessica Sheng, Ori Green, Alvaro A. Ordonez, Mitchell L. Turner, Laura J. Keller, Sanjay K. Jain, Doron Shabat, Matthew Bogyo

**Affiliations:** Department of Pathology, Stanford University School of Medicine, Stanford, CA, USA; School of Chemistry, Raymond and Beverly Sackler Faculty of Exact Sciences, Tel Aviv University, Tel Aviv, Israel; Center for Infection and Inflammation Imaging Research, Johns Hopkins University School of Medicine, Baltimore, MD, USA; Center for Tuberculosis Research, Johns Hopkins University School of Medicine, Baltimore, MD, USA; Department of Pediatrics, Johns Hopkins University School of Medicine, Baltimore, MD, USA; Department of Chemical and Systems Biology, Stanford University School of Medicine, Stanford, CA, USA; Department of Microbiology and Immunology, Stanford University School of Medicine, Stanford, CA, USA

## Abstract

Tuberculosis (TB) is a top-ten cause of death worldwide. Successful treatment is often limited by insufficient diagnostic capabilities, especially at the point of care in low-resource settings. The ideal diagnostic must be fast, cheap, and require minimal clinical resources while providing high sensitivity, selectivity, and the ability to differentiate live from dead bacteria. We describe here the development of a Fast, Luminescent, and Affordable Sensor of Hip1 (FLASH) for the diagnosis and monitoring of drug sensitivity of *Mycobacterium tuberculosis* (*Mtb*). FLASH is a selective chemiluminescent substrate for the *Mtb* protease Hip1 that when processed, produces visible light that can be measured with a high signal to noise ratio using inexpensive sensors. FLASH is sensitive to fmol of recombinant Hip1 enzyme *in vitro* and can detect as few as thousands of *Mtb* cells in culture or in human sputum samples within minutes. The probe is highly selective for *Mtb* compared to other non-tuberculous mycobacteria and can distinguish live from dead cells. Importantly, FLASH can be used to measure antibiotic killing of *Mtb* in culture with greatly accelerated timelines compared to traditional protocols. Overall, FLASH has the potential to enhance both TB diagnostics and drug resistance monitoring in resource-limited settings.

**One Sentence Summary:** A luminescent probe enables sensitive detection of *Mycobacterium tuberculosis* for diagnostics, treatment monitoring, and drug susceptibility testing.

## Introduction

Tuberculosis (TB) is a top-ten cause of death worldwide, with an estimated 10 million new cases leading to nearly 1.5 million deaths yearly. Many of these deaths could be prevented by improving access to diagnostics and therapeutics, especially in low-resource countries that are disproportionally affected. This fact is exemplified by the disparity in disease burdens which range from five or fewer cases per 100,000 in the United States and Europe to more than 500 cases per 100,000 in some countries in Southern Africa and Southeastern Asia. Exacerbating this issue is the extraordinary number of cases that go undiagnosed. The World Health Organization estimates that 3 million cases (30% of all new cases) went unreported in 2018 (*1*). Undiagnosed cases cannot be treated, nor can their spread be mitigated, leading to poor patient outcomes and increased infections. Additionally, diagnoses that take longer than a single clinical visit require the patient to return to receive the test result and begin antibiotic therapy. Diagnostic delay leads to further delays in treatment (*2*). Thus, one of the key pillars of the strategy for eradicating TB is increasing diagnostic capabilities. Rapid, simple, and accurate diagnosis of TB remains a challenge in resource-poor settings.

Current diagnostics are limited by their speed, sensitivity, cost, and their ability to differentiate live from dead bacteria. Culture of *Mycobacterium tuberculosis* (*Mtb*), the causative agent of TB, in liquid or solid media is the gold standard for TB diagnosis. Culture methods offer the highest sensitivity and specificity but are expensive and slow, with conventional culture methods taking up to 8 weeks. In decentralized, resource-poor settings, the recommended diagnostic methods are sputum smear microscopy or the GeneXpert MTB/RIF assay (*3*). Sputum smear microscopy is relatively simple and inexpensive. However, the sensitivity of sputum smear microscopy is dependent on the sputum processing method and the experience of the user, and thus varies widely between 0.32 and 0.97 (*4*), leading to false negatives. The Xpert MTB/RIF assay, developed for the GeneXpert platform, is a nucleic acid amplification test (NAAT) that uses PCR to detect *Mtb* and mutations that confer resistance to rifampicin in under 2 hours, and has sensitivity greater than 0.86 (*5–7*). However, infrastructure requirements, such as continuous electrical supply and trained personnel, limit the implementation of the GeneXpert instrument in peripheral health clinics, and the cost of each Xpert MTB/RIF test in addition to the capital cost for the GeneXpert System is too high for widespread use of this diagnostic method. Furthermore, NAATs like GeneXpert are susceptible to false positives when evaluating disease progression or treatment outcomes because they can amplify bacterial DNA from dead bacteria following antibiotic treatment. Therefore, there is a need for rapid, affordable, point of care diagnostics for TB that offer high sensitivity and selectivity. Ideally, new diagnostic methods should be able to detect small numbers of bacilli and be specific for *Mtb* as opposed to other bacterial species present in the sputum, airway, and oral sites.

The increasing prevalence of multidrug-resistant TB (MDR-TB) also leads to poor health outcomes. In 2018, there were half a million new cases of rifampicin (RIF)-resistant TB (*1*). Early identification of resistance is important for developing proper antibiotic regimens for patients, yet drug susceptibility testing (DST) to determine resistance in clinical isolates can take from 10 days in liquid medium to 28-42 days on solid medium. Molecular tests such as Xpert MTB/RIF provide a rapid method of identifying the presence of mutations that confer resistance, but the design of these tests requires genetic information for each mutation. The development of new tests to detect new resistance mutations or resistance to novel drugs thus requires substantial investment in time and resources. Therefore, new rapid, inexpensive, and comprehensive methods for phenotypic DST have the potential to be transformative to clinical testing of *Mtb* infections.

The development of new diagnostics and methods for DST has attracted substantial attention (*8, 9*). Promising diagnostics under investigation include those that use patient samples other than sputum such as RNA measurements from blood samples (*10*), detection of the *Mtb* cell wall component lipoarabinomannan in patient urine (*11*), and identification of volatile compounds from patient breath (*12, 13*). New fluorescent probes have been developed to report on the activity of mycobacterial enzymes. For example, probes have been developed that target mycobacterial esterases (*14*), sulfatases (*15*), and trehalose mycolyltransesterases (*16, 17*), but these probes are non-specific since they also label nontuberculous mycobacteria (NTMs) that may also be present in sputum. Probes activated by the *Mtb* β-lactamase BlaC have been shown to specifically label *Mtb* in patient sputum samples (*18–20*). These enzyme-based tests offer an improvement in specificity compared to sputum smear microscopy; however, their reliance on fluorescence measurements necessitates imaging instruments that may not be practical for point of care use.

Luminescent probes serve as a promising alternative to fluorescence counterparts. Unlike fluorescence, detection of luminescent signal does not require an excitation source nor advanced optics and can be achieved by a simple, inexpensive luminometer. Background signal is substantially lower because the components of biological samples that often produce undesired fluorescent signals do not spontaneously generate light. Recent advancements in the development of chemiluminescent reporters have made it possible to design aqueous soluble and stable probes that generate light only upon enzymatic cleavage (*21–23*). This approach has been used to create chemiluminescent sensors for β-galactosidase (*24*), cathepsin B (*25*), *Salmonella sp*. esterases and *Listeria monocytogenes* phosphatidylinositol-specific phospholipase C (*26*), and carbapenemase activity in bacteria (*27*). The latter three probes exhibited rapid and sensitive detection of bacteria in culture.

To address diagnostic needs for TB, we have developed a luminescence-based probe that overcomes many of the limitations faced by existing diagnostics and clinical methods for DST. We sought to combine the sensitivity and selectivity of an enzyme-based probe with the ease of detection and low background of a chemiluminescent output. As an enzymatic target we chose the *Mtb* <u>h</u>ydrolase important for pathogenesis 1 (Hip1, Rv2224c). Hip1 is a cell-envelope-associated serine protease that is essential for *Mtb* virulence (*28*) and its survival in macrophages (*29*). Hip1 cleaves the *Mtb* protein GroEL2, contributing to the suppression of early macrophage proinflammatory responses (*30–32*). Together these characteristics make Hip1 an attractive target: (i) its presence on the cell surface makes it highly accessible to small-molecule probes, (ii) its importance for pathogenesis suggests that it will be expressed during infection, (iii) the human genome does not encode a homolog of Hip1, and (iv) prior work from our group has identified an amino acid recognition sequence that is specifically cleaved by Hip1 (*33*). Here we describe a new diagnostic probe for detecting *Mtb*: Fast Luminescent Affordable Sensor of Hip1 (FLASH). FLASH quantitatively reports on the presence of active Hip1 and can quantify and detect as few as 4,000 *Mtb* cells in a one-hour measurement. In human sputum samples, FLASH can detect *Mtb* at concentrations typically found in clinical specimens. Importantly, FLASH also differentiates live from dead bacteria and thus can be used to determine drug susceptibility of clinical isolates using a greatly accelerated and simplified workflow compared to current culture-based methods. Together these data show that FLASH is a promising candidate for rapid TB diagnostics in point of care clinics and clinical DST.

## Results

### Design of a chemiluminescent substrate probe for *Mtb* Hip1

To generate a probe that produces light upon cleavage by Hip1, we combined a selective tetrapeptide Hip1 substrate with a *p*-amino-benzyl-alchohol (PABA) self-eliminating linker and a phenoxy-dioxetane luminophore (Figure 1A). The tetrapeptide sequence was previously optimized for Hip1 cleavage and offers high selectivity for Hip1 compared to other enzymes from *Mtb* or the host (*33*). Upon enzymatic cleavage, the aniline linker undergoes spontaneous elimination, releasing the activated phenoxy-dioxetane luminophore. Subsequent chemiexcitation and decay processes result in the spontaneous generation of light. Live *Mtb* express active Hip1 which processes the probe to produce light (Figure 1B). Emitted light is measured with a sensitive luminometer and integrated over time to yield a total luminescent signal (Figure 1C).

**Figure 1.**
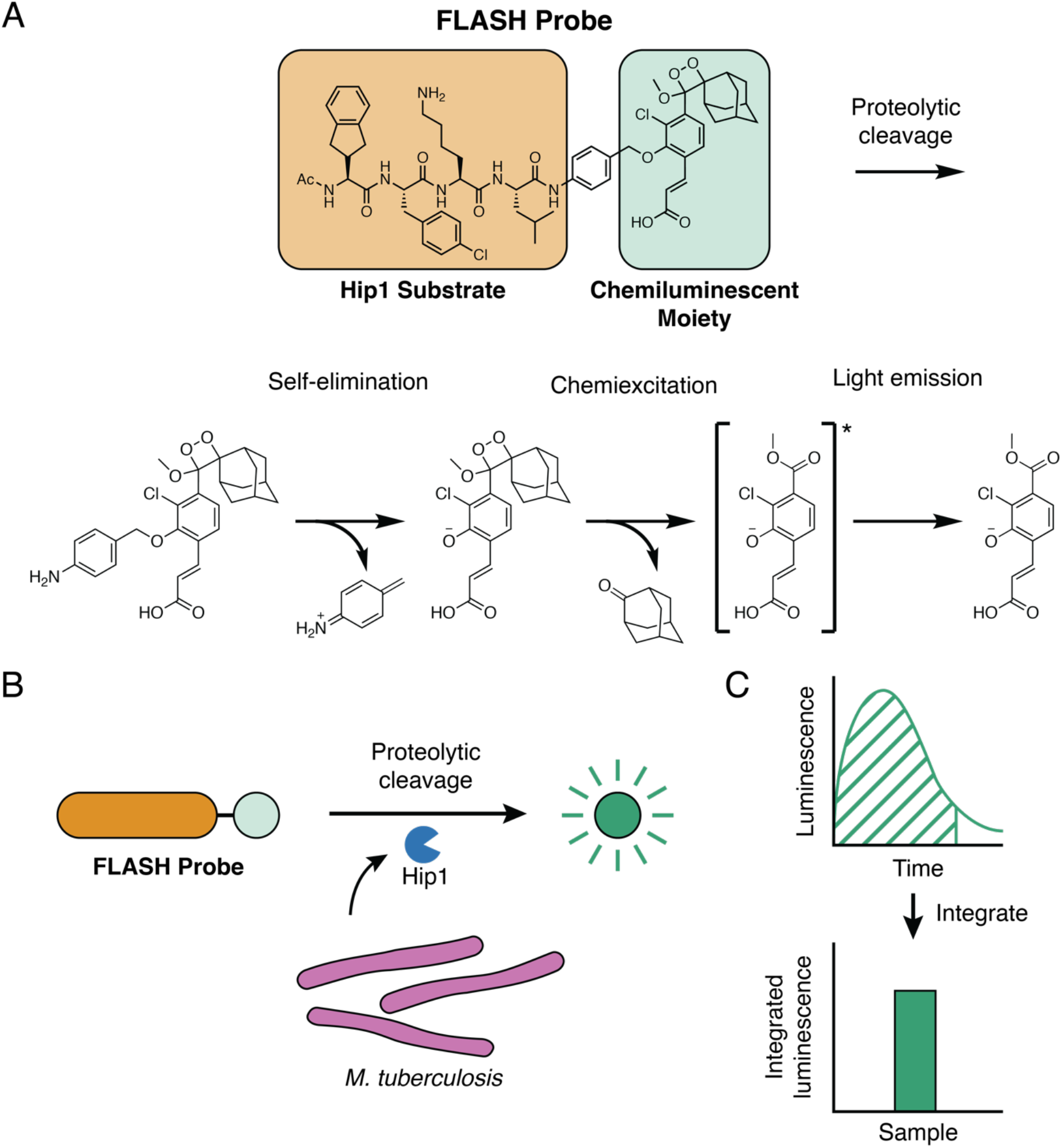
Fast Luminescent Affordable Sensor of Hip1 (FLASH). (A) Following proteolytic cleavage of the FLASH probe by *Mtb* Hip1, self-elimination and chemiexcitation steps ultimately lead to light emission. (B) *Mtb* produces Hip1 protease which cleaves the FLASH probe, producing light. (C) Light produced by probe cleavage is measured over time. Total light output in a given time period (dashed area) is summed to yield integrated luminescence.

We first sought to test that our previously reported fluorogenic substrate probe for Hip1 could be converted to a luminescent reporter. To evaluate Hip1 activity toward the FLASH probe, we titrated recombinantly expressed enzyme and measured luminescent signal with a microplate reader. Light was detected immediately upon enzyme addition and the magnitude of the signal depended on the amount of enzyme added (Figure 2A). Integration of luminescence yielded typical substrate processing curves (Figure 2B). Incubation of Hip1 with a negative control probe containing D-amino acids (D-FLASH, Figure S1) yielded no luminescent signal and provided a measure of background signal resulting from spontaneous release of the reporter due to probe instability (Figure S2). To evaluate the sensitivity of the probe, we compared the integrated FLASH luminescence after 60 min of measurement for each concentration of enzyme to the control samples that lacked enzyme. Samples containing as little as 2.5 fmol enzyme (50 pM) produced a signal that was significantly above the background signal (Figure 2C). Integrated luminescence correlated linearly with total enzyme concentration (r=0.99), indicating that FLASH is both highly sensitive and quantitative for detection of Hip1. Kinetic analysis of titrated probe yielded a k_cat_/K_M_ of 7.5 x 10^4^ M^-1^ s^-1^ (Figure 2D). To verify that the FLASH probe responds only to active enzyme, we incubated Hip1 with the Hip1 inhibitor CSL157 (Figure S1) (*33*) prior to the addition of the FLASH probe. We observed a dose-dependent reduction in luminescence with a calculated IC_50_ of 270 nM (Figure 2E). Results obtained from the recombinant enzyme show that the FLASH probe is a sensitive, quantitative measure of active enzyme. For all FLASH experiments, raw luminescence data (as shown in Fig. 2A) were integrated to yield total luminescence (as shown in Fig. 2C).

**Figure 2.**
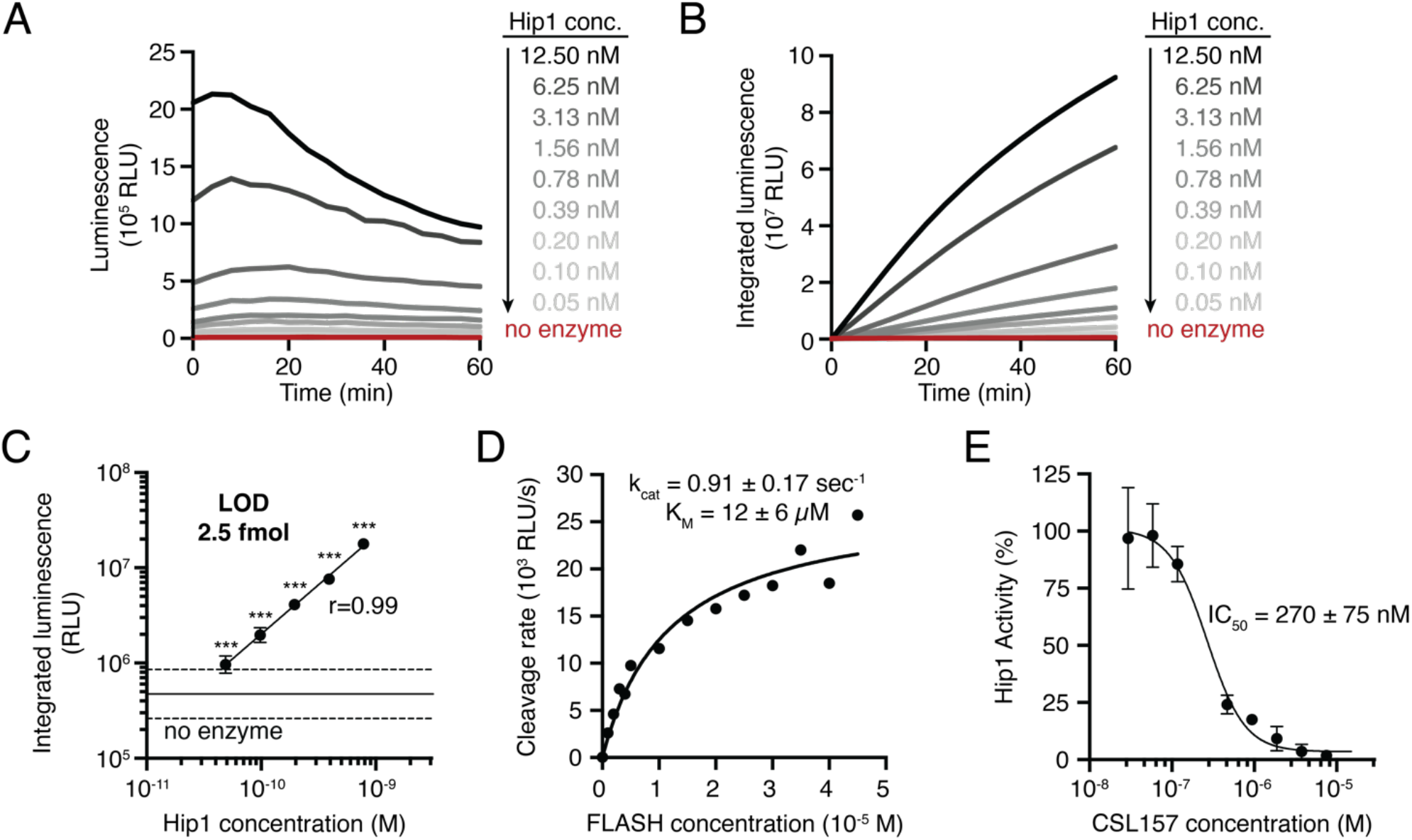
FLASH is a sensitive probe for *Mtb* Hip1 activity. (A) Light emitted by the FLASH probe upon incubation with various concentrations of *Mtb* Hip1. (B) Time course of integrated luminescence from (A). (C) Total integrated luminescence after 1 h incubation with *Mtb* Hip1 (mean ± s.d., n=3). Horizontal lines show the mean (solid) ± 3 s.d. (dashed) of control samples lacking enzyme. Each enzyme concentration was compared to the control samples via one-way ANOVA with Dunnett’s test (***, p<0.001). Best fit line shows linear regression to the log-transformed data. (D) Kinetic analysis of the FLASH probe. Data were fit to the Michaelis-Menten equation to yield kinetic parameters. (E) Inhibition of *Mtb* Hip1 by CSL157 (mean ± s.d., n=3). *Mtb* Hip1 was pre-incubated with inhibitor for 30 min at 37 °C before addition of FLASH probe. Data were fit to a four-parameter logistic equation to yield IC_50_. All parameters are reported as the 95% confidence interval.

### FLASH is a quantitative sensor of live *Mtb*

To determine whether the FLASH probe can detect live *Mtb*, we added probe to bacterial cultures and measured luminescence. To optimize assay conditions for bacterial detection, the signal to noise ratio (SNR) was calculated for different integration times by dividing the integrated signal for cultures with and without *Mtb* cells (Figure S3A). Sixty minutes of integration time yielded a SNR of greater than 15, and we used this measurement time for all subsequent experiments. Luminescent signal was reduced when cultures were preincubated with Hip1 inhibitor (Figure S3B), indicating that, as observed for experiments with the recombinant enzyme, probe cleavage in the presence of cells is dependent on active Hip1.

A key metric for evaluating FLASH as a potential diagnostic is the ability for the probe to detect the small numbers of bacteria usually found in sputum samples. To quantify the limit of detection of FLASH for *Mtb*, bacterial cultures were serially diluted into either culture medium or processed human sputum prior to adding the probe (Figure 3A). In all cases, absorbance (OD_600_) was used as a proxy for cell number. The conversion factor between absorbance and cell count was obtained by CFU plating (OD_600_ 1 = 3 x 10^8^ CFU/mL). The experiment was performed with two strains of *Mtb*: the laboratory strain, H37Rv, measured in a biosafety-level-3 (BSL3) facility and the attenuated strain, mc^2^6020 (*34*), measured in a BSL2 facility. For both strains, luminescent signal was linearly correlated with cell number (Figure 3B-C). Limits of detection (LOD) were calculated by extrapolating best fit lines to estimate the cell number for which luminescent signal would equal three times the standard deviation of the negative controls lacking cells. LODs for H37Rv and for mc^2^6020 were 23,000 and 4,000 cells, respectively. We attribute the higher LOD in H37Rv to the less sensitive microplate reader available for use in the BSL3. SNRs were higher at all integration times for the more sensitive BSL2 microplate reader (Figure S3A) and there was not a significant difference between integrated luminescence for H37Rv and for mc^2^6020 when both were measured on the BSL3 microplate reader (Figure S3C).

**Figure 3.**
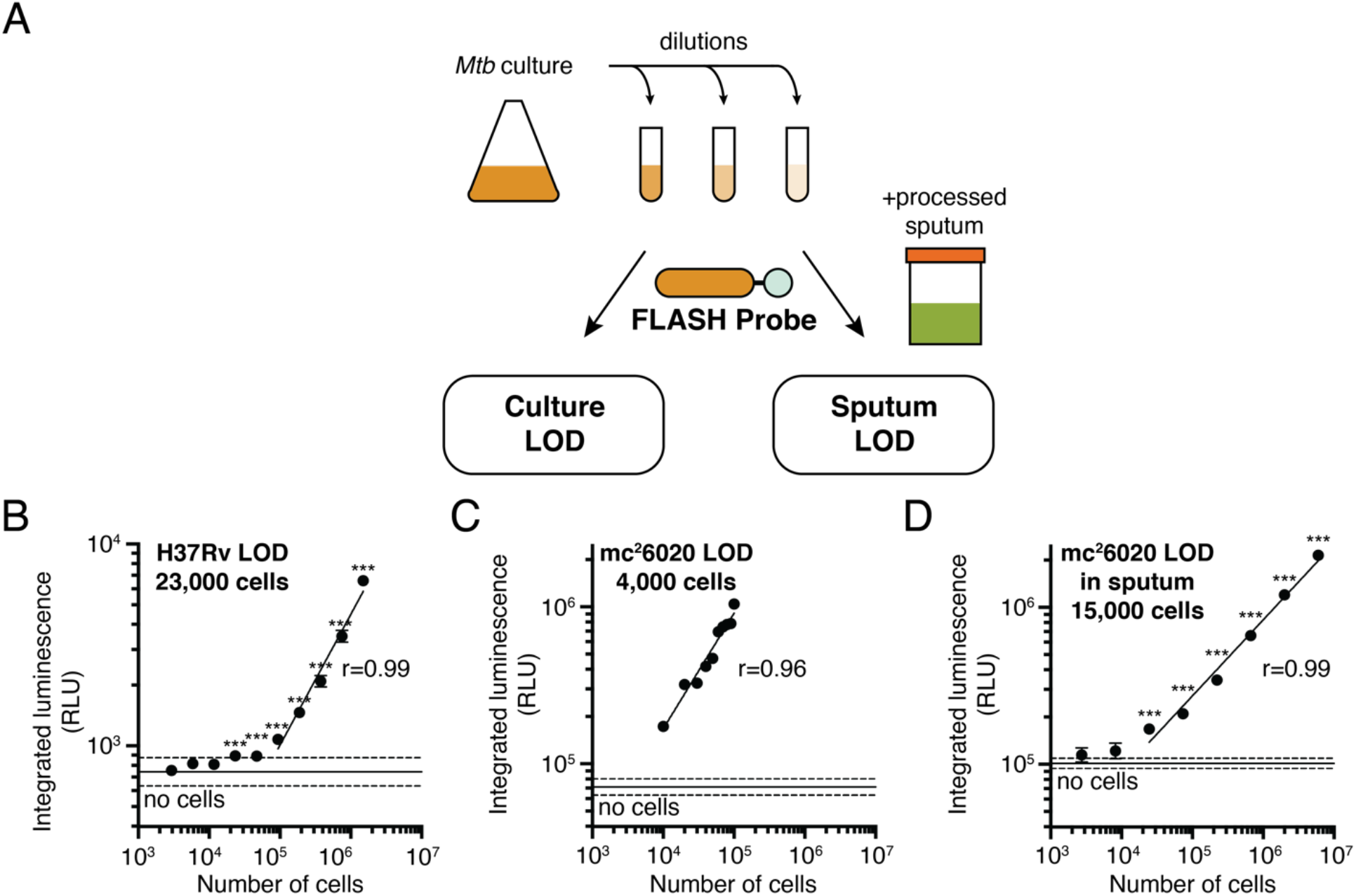
FLASH is a quantitative probe for *Mtb* cells. (A) To determine limit of detection, cultures of *Mtb* were serially diluted into medium or into processed human sputum and then incubated with the FLASH probe. (B-D) Total integrated luminescence after 1 h incubation of FLASH probe with (B) *Mtb* H37Rv in 7H9 medium, (C) mc^2^6020 in 7H9 medium, and (D) mc^2^6020 in processed human sputum (mean ± s.d., n=3). Horizontal lines show the mean (solid) ± 3 s.d. (dashed) of control samples lacking cells. For all experiments, each sample was compared to the no-bacteria control via one-way ANOVA with Dunnett’s test (***, p<0.001; for (C) all comparisons yielded p<0.001). Best fit lines show linear regressions to the log-transformed data (excluding cell concentrations that are not significantly different from the control). Limits of detection were calculated by determining the cell number for which the best fit line intercepts the mean + 3 s.d. of the control samples.

To evaluate whether the high sensitivity of FLASH observed for bacteria in culture is likely to translate to diagnostic detection in clinical samples, we spiked mc^2^6020 grown in culture into pooled human sputum collected from patients who had tested negative for TB. The sputum was processed using the standard NALC/NaOH protocol (*35*) at the time of collection. Sputum processing was optimized by centrifuging sputum samples and resuspending the pellet in PBS to achieve neutral pH. This step resulted in an 8-fold increase in signal (Figure S3D). Using the centrifugation and resuspension protocol, the LOD for mc^2^6020 in processed sputum was 15,000 cells, again with a linear correlation between cell number and luminescent signal. The higher LOD measured in sputum compared to culture is consistent with a slight loss of signal observed after the centrifugation step. Signal generated from bacteria in sputum was lower than the same amount of bacteria added directly to buffer, but this loss of signal was recapitulated by centrifugation of bacteria in culture (Figure S3D), suggesting that it was the centrifugation step and not the sputum itself that reduced the signal. Although Hip1 is reported to be surface-associated, one explanation for the decreased signal could be that some enzyme is released into the culture medium. To test this, we centrifuged bacterial cells and compared the luminescent signal among the original culture, the supernatant, and the resuspended pellet. Indeed, a substantial fraction of the culture signal was present in the supernatant (40%), with the remainder found in the pellet (55%; Figure S3E), suggesting that either free Hip1 enzyme or some fraction of cells with Hip1 activity are found in the supernatant following the centrifugation step.

The limit of detection for cells in processed sputum is encouraging for the ability of FLASH to diagnose TB. The concentration of *Mtb* bacilli in sputum ranges broadly across patients, according to disease severity, and throughout the course of antibiotic treatment. Measurements based on colony forming units (CFU) on solid culture range from 10^1^ to 10^7^ CFU/mL sputum but data from a variety of studies agree on average concentrations between 10^5^ to 10^6^ CFU/mL (*36–38*). With the concentration step routinely used in sputum processing, a typical 1 mL sample of sputum is expected to have at least an order of magnitude more cells than our limit of detection.

### FLASH is specific for *Mtb*

A key feature of a diagnostic for *Mtb* is that the method does not provide false positives due to other bacteria or enzymes that may be present in clinical sputum samples. Potential sources of false positives include host enzymes present in the saliva or sputum, constituent organisms of the healthy microbiota, or NTMs. The human genome does not encode a Hip1 homolog and the low background signal observed in processed human sputum (Figure 3D) suggests that enzymes from the host or the microbiome do not activate the probe or do not survive the processing protocol. We used phylogenetic analysis of proteins with sequence similarity to *Mtb* Hip1 to identify organisms that may lead to false positives. We searched for proteins with sequence similarity to Hip1 in the Human Microbiome Project databases of oral and airway microbiota (*39, 40*), a set of NTMs, and other pathogens known to infect the human lung (*Staphylococcus aureus* and *Klebsiella pneumoniae*). Putative Hip1 homologs from these organisms clustered into two groups: a set of closely related enzymes with at least 65% similarity found in NTMs and a more divergent set found in other bacteria (Figure 4A, Figure S4A). The latter group of enzymes shared higher sequence similarity with the *Mtb* α/β hydrolase Rv2223c than with Hip1. This is notable because our prior work showed that the fluorogenic precursor to the FLASH probe is not processed by a *hip1* knockout strain of *Mtb* (*29, 33*), suggesting that the peptide sequence is not a substrate for Rv2223c.

**Figure 4.**
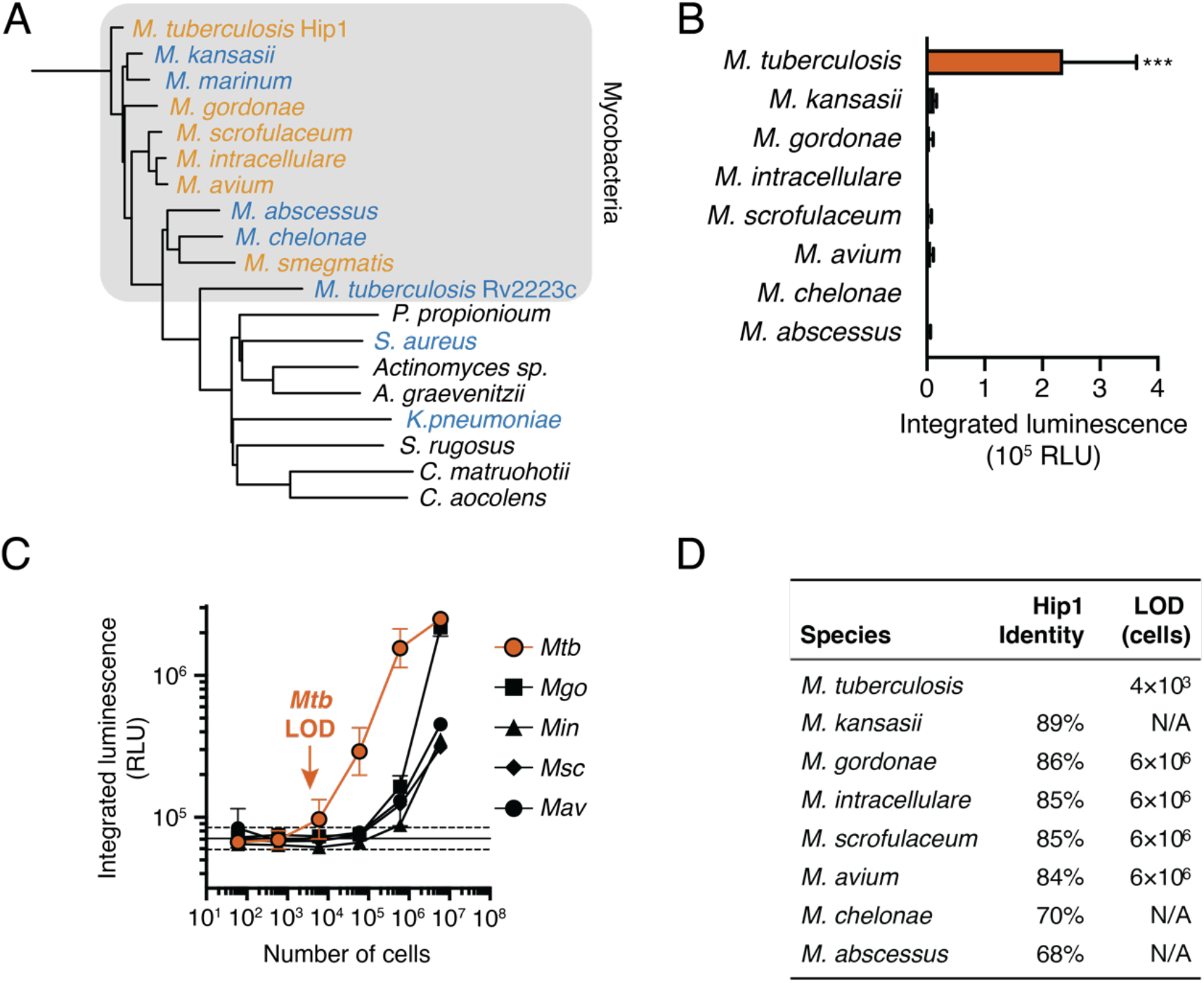
FLASH is selective for *Mtb*. (A) Phylogenetic tree of potential *Mtb* Hip1 homologs found in NTMs, other lung pathogens, and commensal members of the human airway and oral microbiomes. Also included is Rv2223c, an uncharacterized peptidase encoded by *Mtb* with sequence similarity to Hip1. Bacteria are colored by their ability to process FLASH at high cell densities (orange, active; blue, inactive; black, not tested). (B) FLASH signal for 6×10^4^ cells of each NTM in 7H9 medium (mean ± s.d., n=3). Each sample was compared to the no-bacteria control via one-way ANOVA with Dunnett’s test (***, p<0.001). (C) FLASH signal for *M. tuberculosis* (*Mtb*), *M. gordonae* (*Mgo*), *M. intracellulare* (*Min*), *M. scrofulaceum* (*Msc*), and *M. avium* (*Mav*) at the indicated cell number (mean ± s.d., n=3). (D) Sequence similarity of potential homologs to *Mtb* Hip1 and limits of detection calculated from (C) for each NTM.

To evaluate probe specificity, we measured the FLASH signal in bacterial cultures of NTMs and other common pathogens. Three commonly used laboratory strains of *Mtb* (H37Rv, CDC1551, and Erdman) as well as the disease-causing *Mycobacterium bovis* all yielded signal substantially above background (Figure S4B). At high cell densities (3×10^8^ CFU/mL; 1.2×10^7^ cells) clinical isolates of *Mycobacterium gordonae, Mycobacterium intracellulare* (*Min*), *Mycobacterium scrofulaceum* (*Msc*), *Mycobacterium avium* (*Mav*), and the laboratory strain of *Mycobacterium smegmatis* (*Msm*) yielded luminescent signal above background, but substantially lower than for *Mtb* (Figure S4C-D). Furthermore, at a 200-fold lower cell density that is more representative of bacterial burdens in TB sputum (1.5×10^6^ CFU/mL; 6×10^4^ cells), only cultures of *Mtb* yielded luminescent signal significantly greater than background (Figure 4B). To compare activation of the FLASH probe among NTMs, we calculated LODs for each species that showed some activation (Figure 4C). All LODs were 1,000-fold higher than for *Mtb* (Figure 4D), indicating that FLASH is highly selective and is unlikely to give false positive signals in response to NTMs.

### FLASH enables rapid drug susceptibility testing

The ability to distinguish live from dead *M. tuberculosis* is another key feature of a TB diagnostic. Similarly, the ability to detect antibiotic killing of isolates is critical in clinical microbiology, especially for TB for which drug resistance is a growing global threat. We hypothesized that FLASH could be used to distinguish live from dead bacteria. Initial experiments showed that the luminescent signal was greatly reduced in heat-killed cultures or following treatment of cultures with the antibiotic rifampicin (RIF, Figure S5A). This observation suggests that cell death results in decreased levels of active Hip1, presumably due to protein degradation or instability combined with the cessation of new protein synthesis. To further evaluate the potential for FLASH to report on cell viability we treated *Mtb* cultures with RIF over the course of nine days. Cultures were sampled throughout the treatment course and cell viability was measured using a 24 h treatment with resazurin (CellTiter-Blue, CTB). The same cultures were incubated with the FLASH probe for 1 h (Figure 5A). RIF itself absorbs light in the visible range (absorption maximum at 475 nm (*41*)), but at the concentrations tested, it had no effect on luminescent signal from the dioxetane luminophore (Figure S5B). We observed a high correlation (r=0.95) between FLASH luminescence and CTB fluorescence across the range of RIF concentrations (Figure 5B) indicating that FLASH is a quantitative indicator of cell viability. We generated dose-response curves by comparing FLASH signal to RIF concentration (Figure 5C). At early time points (days one and three), there was no difference between high and low concentrations of RIF, but at later time points we observed the expected dose response curves, with a low FLASH signal in the cultures treated with RIF concentrations that prevent growth. At day seven, the dose response curve generated using FLASH yielded an IC_50_ of 36 ± 21 nM (Figure 5D), matching the EC_50_ value calculated from CTB measurements of the same cultures (Figure 5E).

**Figure 5.**
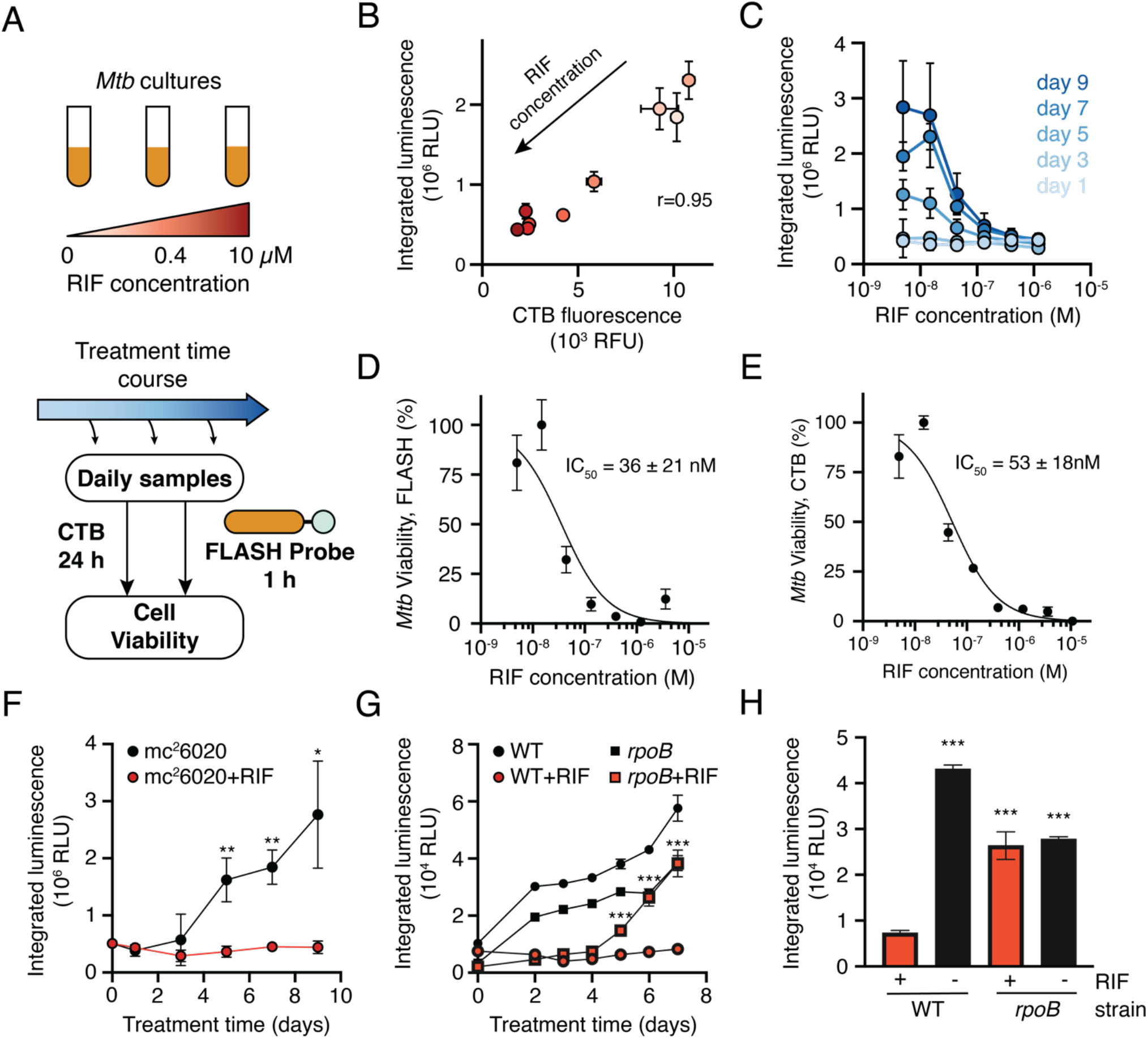
FLASH provides a quantitative measure *Mtb* viability. (A) *Mtb* cultures were treated with RIF for up to nine days. Samples were removed throughout the treatment period and incubated with FLASH for 1 h, or with CellTiter-Blue (CTB) for 24 h. (B) FLASH and CTB measurements for cultures treated for 7 d with RIF (mean ± s.d., n=3). Marker colors correspond to RIF concentrations shown in (A). (C) FLASH signal dependence on RIF concentration for each day (mean ± s.d., n=3). (D-E) Dose response for killing by RIF as measured by the FLASH probe (D) or CTB (E) (mean ± s.d., n=3). Data were normalized to DMSO (100% viability) and 10 μM RIF (0% viability) and fit to a two-parameter logistic function. IC_50_ values are reported as 95% confidence intervals. (F-G) Time course of (F) mc^2^6020 or (G) H37Rv (WT) *Mtb* and RpoB H526D mutant *Mtb* (*rpoB*) treated with DMSO (black) or the critical concentration of RIF (1.2 μM, red). For each day, the RIF- and DMSO-treated conditions were compared via an independent t-test (n=3; ***, p<0.001; **, p<0.01; *, p<0.05). (H) Luminescent signal from H37Rv (WT) or *rpoB* after 6 d of culture in the presence or absence of RIF. Samples are compared to the WT *Mtb* strain treated with RIF via one-way ANOVA with Dunnett’s test (***, p<0.001).

To evaluate the potential for FLASH as a tool for drug-susceptibility testing (DST), we treated cultures of mc^2^6020 with the concentration of RIF used clinically to determine drug susceptibility (1.2 μM; 1 mg/L) and compared the FLASH signal between treated and untreated cultures throughout the treatment period (Figure 5F). We observed a significantly lower signal in the treated samples as soon as five days after treatment. For clinical use, a DST protocol must be able to distinguish susceptible and resistant strains. To test for its ability to identify drug resistance, we repeated the DST experiment and compared a RIF-susceptible (H37Rv) to a RIF-resistant strain (RpoB H 526D mutant, *rpoB*) (Figure 5G). Luminescence increased over time for the untreated cultures and for the resistant strain treated with RIF. As observed before, there was no change in signal for the susceptible strain treated with RIF. After six days of treatment, signal for the resistant strain treated with RIF had increased substantially and all samples yielded significantly higher signal than the susceptible strain treated with RIF (Figure 5H). Similar results were obtained when comparing the RIF-susceptible strain CDC1551 to the RIF-resistant strain (Figure S5C). These results show that FLASH can be used to differentiate susceptible from resistant strains of *Mtb* after five to six days of culture with antibiotic—substantially faster than the weeks to months generally required for culture-based DST, and faster than newer methods for DST such as microscopic-observation drug-susceptibility (8 days) (*42*) and Sensititre MycoTB Plates (10-14 days) (*43*).

## Discussion

Global health initiatives including The World Health Organization (*1*) and the Stop TB Partnership (*44*) have highlighted rapid point-of-care diagnostics and rapid DST as critical developments needed to reach the goal of reduced TB cases and better patient outcomes. FLASH has the potential to address these needs by providing a fast, sensitive, and selective tool for the detection of live *Mtb* in culture or sputum samples. The labeling protocol is simple, requires minimal training, and does not require microscopy or other sophisticated laboratory instrumentation. Our experiments with processed sputum show that it is compatible with the standard NALC/NaOH protocol used for decontaminating sputum samples. Because FLASH is selective for *Mtb* compared to other organisms that may be present in sputum samples, the readout is not sensitive to potential contaminants and this decontamination step may be unnecessary. We expect that further optimization of sputum processing and concentration of bacteria will reduce the loss of signal due to centrifugation, decreasing the limit of detection in sputum samples.

The ability of FLASH to differentiate live from dead bacteria also offers the potential for its use as a rapid readout for monitoring TB treatment. Despite their high sensitivity, NAATs like Xpert are problematic when used for diagnosis following treatment because they can amplify DNA left over from dead bacteria. These false positives can persist for years after treatment (*45, 46*) making it difficult for NAATs to accurately track decreasing disease burden during antibiotic therapy (*47*). In contrast, FLASH does not respond to dead bacteria and should be able to track the decrease in bacterial burden throughout a course of antibiotic treatment. In addition, the short time to result for FLASH compared to culture-based method should enable rapid assessment of treatment outcomes, which is important for identifying and adapting to treatment failures due to antibiotic resistance.

In this study we tested three different plate readers to evaluate the sensitivity and selectivity of FLASH and we note that differences in instrumentation lead to differences in the limits of detection and signal to noise ratios (Figure 3B-C, Figure S3A). For point of care diagnostics using FLASH it will be critical to use a sensitive, inexpensive, and low power luminometer. A number of devices that meet these criteria have been evaluated for other purposes including hand-held, battery-operated luminometers (*48*) and adapters for use with smartphone cameras (*49, 50*). Additional studies will be required to evaluate the ability of FLASH to (i) sensitively and specifically detect TB at the point of care, (ii) track disease progression throughout the course of antibiotic treatment, and (iii) reliably determine the antibiotic susceptibility of clinical isolates more quickly than existing DST approaches.

This study along with other examples of luminescent enzyme probes for pathogen detection (*26*) highlight the versatility and adaptability of turn-on dioxetane luminophores. Key advantages of luminescent measurements are the low background signal in biological samples and the simplicity of detection. The design of these probes is straightforward—any enzymatic or chemical unmasking event can lead to light emission. Proteases offer attractive targets for such probes because of the flexibility to design peptidic substrates with great selectivity. By carefully choosing the appropriate enzyme target and cognate substrate, we expect luminescent turn-on probes will serve as effective point-of-care diagnostics for other infectious diseases in addition to TB.

## Materials and Methods

### Study Design

This study comprises controlled laboratory experiments designed to determine the sensitivity and selectivity of the FLASH probe in detecting and quantifying Hip1 enzyme activity and live *M. tuberculosis*. Units of investigation included recombinant Hip1 enzyme, cultures of *M. tuberculosis* or other bacteria and de-identified human sputum samples. Unless otherwise indicated, all experiments were performed with biological triplicates. Measurements of *M. tuberculosis* detection were performed by four researchers at three sites (Stanford University BSL2, Stanford University BSL3, and Johns Hopkins University BSL3 facilities) using three different plate readers. No data were excluded from the analyses. Experiments were not blinded.

### Chemical Synthesis

Methods for the synthesis and characterization of the FLASH and D-FLASH probes are presented in the Supporting Information.

### Bacterial Culture

*M. tuberculosis* H37Rv, *Mycobacterium marinum* M, and *Mycobacterium smegmatis* mc^2^155 were a gift from Carolyn Bertozzi (Stanford University). *M. tuberculosis* mc^2^6020 was a gift from Niaz Banaei (Stanford University). *M. tuberculosis* Erdman and the CDC1551-derived RpoB H526D rifampicin-resistant mutant (*51*) were obtained from the Center for Tuberculosis Research (Johns Hopkins University). The following clinical isolates of NTMs were a gift from Dr. Nikki Parrish and Derek Armstrong (Department of Pathology, the Johns Hopkins University School of Medicine): *Mycobacterium kansasii, Mycobacterium gordonae, Mycobacterium intracellulare, Mycobacterium scrofulaceum, Mycobacterium avium*, *Mycobacterium chelonae*, and *Mycobacterium abscessus*.

*M. tuberculosis* strains (except mc^2^6020) and all NTMs were cultured in liquid 7H9/OADC medium (4.7 g/L 7H9 powder, 0.2% w/v glycerol, 0.05 % w/v Tween-80, and 10% v/v OADC supplement) or on solid 7H10 agar plates (19 g/L 7H10 powder, 1% w/v glycerol, 10% OADC supplement). *M. tuberculosis* mc^2^6020 was cultured in liquid 7H9/OADC medium supplemented with 24 mg/L pantothenate, 80 mg/L L-lysine, and 0.2% w/v casamino acids or solid 7H9 plates (15 g agar, 4.7 g 7H9 powder, 0.1 % w/v glycerol, 0.2 % w/v casamino acids, 24 mg/L pantothenate, 80 mg/L L-lysine, 10% OADC supplement). OADC supplement contained 0.5 g/L oleic acid, 50 g/L albumin fraction V, 20 g/L dextrose, 40 mg/L catalase, and 8.5 g/L NaCl. Cultures were inoculated from frozen glycerol stocks or from agar plates and cultured at 37 °C with shaking for at least one week. To estimate the number of cells used for each experiment, a conversion factor was calculated by plating serial dilutions of cultures with known OD_600_ values onto agar plates. After 3-5 weeks of growth at 37 °C, individual colonies were counted to determine CFU/mL. Two separate experiments yielded the same factor of OD_600_ 1 = 3×10^8^ CFU/mL. For each experiment, OD_600_ was measured in a spectrophotometer and cultures were diluted to the desired cell density based on the conversion factor.

### FLASH Measurements and Data Analysis

All experiments were performed in triplicate, unless otherwise indicated. All chemiluminescence assays were performed in white, opaque flat-bottom 384- or 96-well plates. Luminescence was measured in different microplate readers, depending on the laboratory location. Measurements of H37Rv, mc^2^6020, and *M. marinum* were obtained on a SpectraMax M3 (Molecular Devices) at 25 °C. Measurements of recombinant Hip1, mc^2^6020, NTMs, and other bacteria were obtained on a Cytation 3 (Biotek) at 37 °C. Measurements of rifampicin susceptibility of H37Rv and RpoB H526D mutant were obtained on a FLUOstar Omega (BMG Labtech) at 37 °C. For all experiments, luminescence measurements began within 5 min after the addition of FLASH probe and were continued for at least 1 h. Measurements were made without an emission filter, using a 1 s integration time. For each sample, luminescence measurements from the first hour were summed to yield integrated luminescence.

### Detection of Hip1 Enzyme Activity

Recombinant Hip1 was purified as previously described (*33*). For each experiment, 40 μL of Hip1 in Hip1 buffer (0.01% Triton X-100 in PBS) was combined with 5 μL of 9x FLASH probe in 1:1 DMSO/Hip1 buffer. To determine the limit of detection, 40 μL of two-fold series dilutions of recombinant Hip1 (final concentrations: 0.05-12.5 nM) were combined with 5 μL of 90 μM FLASH probe (final concentration 10 μM). To determine kinetic parameters of the probe, 40 μL of 3 nM Hip1 in Hip1 buffer were combined with a dilution series of FLASH probe concentrations (final concentrations: 0-50 μM). To measure enzyme inhibition, 37.5 μL of 3 nM Hip1 in Hip1 buffer were combined with 2.5 μL of CSL157 in DMSO (final concentrations: 20 nM-7 μM) and incubated for 30 min at 37 °C before addition of 5 μL of FLASH probe (final concentration: 10 μM). To test D-FLASH, 40 μL of 3 nM Hip1 in Hip1 buffer was combined with 10 μM of D-FLASH probe. All conditions were tested in triplicate.

### Analysis of Hip1 Homologs

The protein sequence of Hip1 (Rv2224c) from *M. tuberculosis* H37Rv was used as the query sequence for BLASTP analysis. The search set comprised whole genome sequences from NTMs, the Human Microbiome Project Reference Genome reference isolates in the airways and oral subsets (https://www.hmpdacc.org/hmp/HMRGD/, retrieved on April 28, 2020), *Staphylococcus aureus*, and *Klebsiella pneumoniae*. Hip1 was queried against the protein sequences by BLASTP (2.9.0+), requiring a minimum e-value of 1e-4. The sequences of Hip1, *M. tuberculosis* Rv2223c, and the closest homolog of Hip1 in NTMs, *S. aureus, K. pneumoniae*, and select reference bacteria from the oral and airway microbiomes were aligned using Clustal Omega (https://www.ebi.ac.uk/Tools/msa/clustalo/) (*52*), and the phylogenetic tree was generated using FigTree (1.4.4).

### Detection of Bacteria in Culture

Cultures were grown until reaching an OD_600_ of 0.4–1.0 and then were diluted in growth medium to reach the desired cell concentration for each condition. To determine the limit of detection, 40 μL of serially diluted cultures were added to a 384-well plate. Heat-killed control samples were prepared by heating 0.5 mL of culture in an O-ring tube at 95 °C for 15 min. Hip1 inhibition in live cells was tested by treating 3×10^9^ CFU/mL H37Rv with 10 μM CSL157 for 1 h at 37 °C before addition of FLASH. Where indicated, cultures were centrifuged for 10 min at 8,000 rcf, the supernatant was removed by pipetting, and the cell pellet was resuspended in PBS. For all experiments, 5 μL of 225 μM FLASH probe in 1:1 DMSO/Hip1 buffer (final concentration 25 μM) was added to 40 μL of bacterial sample. All conditions were tested in triplicate.

### Detection of Bacteria in Sputum

Sputum samples were obtained from the Johns Hopkins Clinical Microbiology Laboratory as per standard of care. Sputum was processed using a standard decontamination and concentration protocol. Sputum was transferred to a 50 mL centrifuge tube, combined with an equal volume Snap n’ Digest (Scientific Device), vortexed, and incubated at 25 °C for 15 min. Samples were neutralized by addition of PBS to a final total volume of 45 mL. Samples were centrifuged at 3200 rcf and the supernatant was removed. Pellets were resuspended in up to 5 mL of PBS. Decontaminated samples from multiple patients were pooled, aliquoted, and frozen at −80 °C until use. Cultures of mc^2^6020 were grown until reaching an OD_600_ of 0.2. Cultures were diluted in growth medium to reach the desired cell concentration for each condition and added to an equal volume of processed human sputum. Samples were neutralized by addition of PBS and then centrifuged for 17 min at 3200 rcf. Supernatant was removed and the pellet was resuspended in 130 μL of PBS. For all experiments, 5 μL of 225 μM FLASH probe in 1:1 DMSO/Hip1 buffer (final concentration 25 μM) was added to 40 μL of bacterial sample. All conditions were tested in triplicate.

### Drug Susceptibility Testing

Cultures of *M. tuberculosis* were grown until reaching an OD_600_ of 0.4–1.0 and diluted to an OD_600_ of 0.2 in growth medium. The diluted culture was aliquoted for each treatment condition into 5-10 mL cultures. Rifampicin was added from 100x stock made up in DMSO to the desired concentration. Cultures were shaken at 37 °C for the duration of the experiment. At each time point, an aliquot of culture was transferred to a 384-well plate. For all experiments, 5 μL of 225 μM FLASH probe in 1:1 DMSO/Hip1 buffer (final concentration 25 μM) was added to 40 μL of bacterial sample. To test for cell viability, 100 μL of each culture was transferred to a 96-well plate, treated with 20 μL of CellTiter-Blue (Promega), and incubated for 24 h at 37 °C. CellTiter-Blue fluorescence was measured with 560nm excitation and 590nm emission. All conditions were tested in triplicate.

## Supplementary Materials

Supporting Methods (53, 54)

Figure S1. Chemical structures.

Figure S2. Activities of Hip1 toward FLASH and D-FLASH.

Figure S3. Comparison of *M. tuberculosis* strains and sputum treatment conditions.

Figure S4. Comparison of FLASH signal from NTMs and other bacteria.

Figure S5. FLASH and CTB measurements of RIF-treated cultures.

## Acknowledgements

We thank Dr. Carolyn Bertozzi and Dr. Douglas Fox (Stanford University) for sharing bacterial strains and for training and use of the Stanford BSL3 facility. We thank Dr. Niaz Banaei (Stanford University) for sharing bacterial strains. We thank Dr. Nikki Parrish and Derek Armstrong (Department of Pathology, the Johns Hopkins University School of Medicine) for sharing bacterial strains and de-identified human sputum samples.

## Author Contributions

**Brett M. Babin:** Conceptualization, Formal Analysis, Investigation, Visualization, Writing – original draft; **Gabriela Fernandez-Cuervo:** Conceptualization, Formal Analysis, Funding Acquisition, Investigation, Resources, Writing – review & editing; **Jessica Sheng:** Investigation, Formal Analysis, Writing – review & editing; **Ori Green**: Investigation, Resources; **Alvaro A. Ordonez**: Conceptualization, Investigation, Funding Acquisition; **Mitchell L. Turner:** Investigation; **Laura J. Keller:** Investigation, Writing – review & editing; **Sanjay K. Jain:** Resources, Funding Acquisition; **Doron Shabat:** Conceptualization, Funding Acquisition, Supervision, Writing – review & editing; **Matthew Bogyo:** Conceptualization, Funding Acquisition, Supervision, Writing – review & editing. **Competing Interests.** None declared. **Data and materials availability.** All data associated with this study are available in the main text or the supplementary materials.

