## Supplementary material for "A chemiluminescent protease probe for rapid, sensitive, and inexpensive detection of live *Mycobacterium tuberculosis*": Supplemetal Figures and Methods

### Supporting Figures

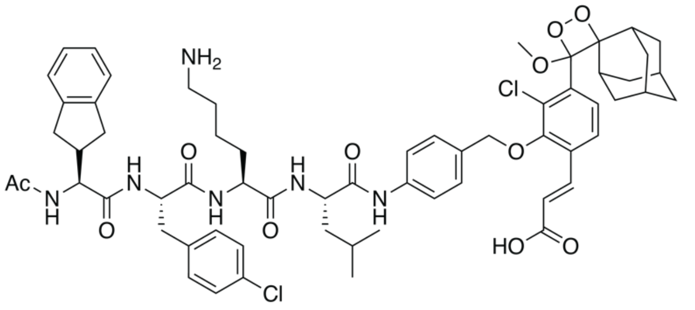

**FLASH Probe**
Ac-Igl-4-Cl-Phe-Lys-Leu

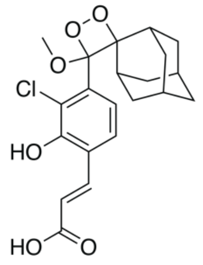

**AquaSpark**

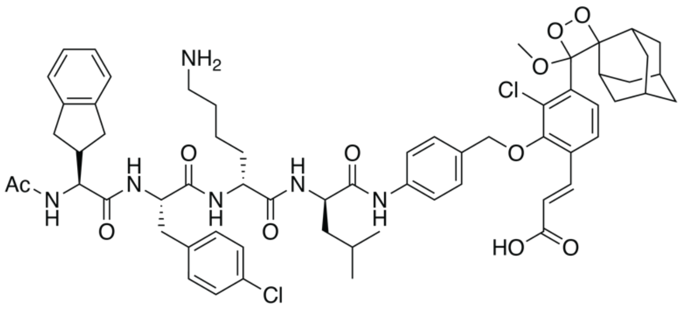

**D-FLASH Probe**
Ac-Igl-4-Cl-Phe-D-Lys-D-Leu

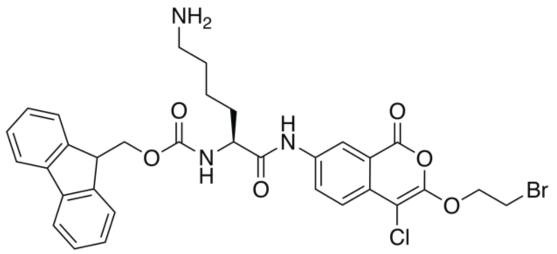

**CSL157**

**Figure S1. Chemical structures.**

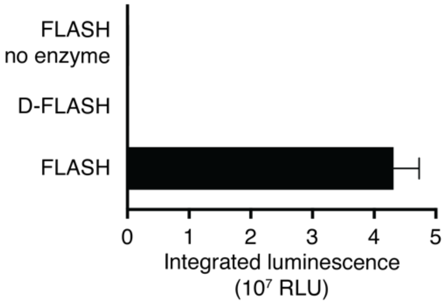


**Figure S2. Activities of Hip1 toward FLASH and D-FLASH.** FLASH and D-FLASH probes (10 μM) were incubated with recombinant 3 nM Hip1. The control sample contains FLASH but no enzyme.

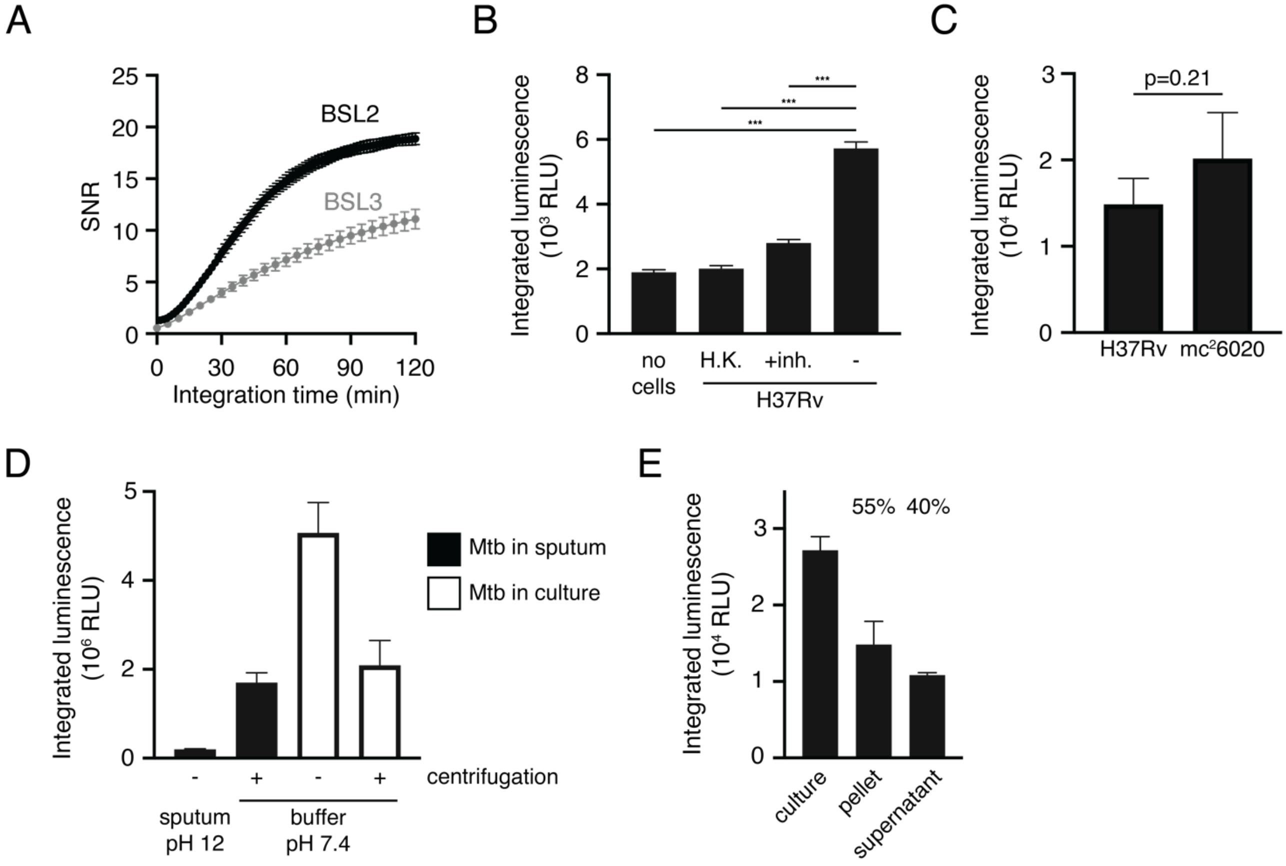


**Figure S3. Comparison of *M. tuberculosis* strains and sputum treatment conditions.** (A) Signal-to-noise ratios (SNR) for different integration times for *M. tuberculosis* mc$^2$6020 measured on the microplate readers in the BSL2 and BSL3 facilities (mean ± s.d., n=3). (B) Cultures of *M. tuberculosis* H37Rv were heat killed (H.K.) or incubated with 10 μM CSL157 for 1 h prior to addition of the FLASH probe and measurement of integrated luminescence (mean ± s.d., n=3). The control sample contains FLASH probe but no cells. Sample means were compared among all conditions via Tukey's test (***, p<0.001). (C) Integrated luminescence for *M. tuberculosis* H37Rv and M. tuberculosis mc$^2$6020 measured on the BSL3 microplate reader (mean ± s.d., n=3). Sample means were compared via t-test. (D) Integrated luminescence for *M. tuberculosis* mc$^2$6020 spiked into processed sputum (black) or from culture (white). Samples were centrifuged and resuspended in neutral pH buffer as indicated (mean ± s.d., n=3). (E) Integrated luminescence for a culture of *M. tuberculosis* mc$^2$6020 before (culture) and after centrifugation and separation into pellet and supernatant fractions (mean ± s.d., n=3). Percentages above the fractions indicate the fraction of signal from the original culture.

A

| Species | Identity | Ser 228 | Asp 463 | His 490 |
|---|---|---|---|---|
| *M. tuberculosis* | | TYLGY**S**YGTRI | VSTTH**D**PATPY | FDGTQ**H**TVVFQ |
| *M. kansasii* | 89% | TYLGY**S**YGTRI | VSTTH**D**PATPY | FNGTQ**H**TVVFQ |
| *M. gordonae* | 86% | TYLGY**S**YGTRI | VSTTH**D**PATPY | YNGTQ**H**TVVFQ |
| *M. intracellulare* | 85% | TYLGY**S**YGTRI | VSTTH**D**PATPY | YDGTQ**H**TVVFQ |
| *M. scrofulaceum* | 85% | TYLGY**S**YGTRI | VSTTH**D**PATPY | YDGTQ**H**TVVFQ |
| *M. avium* | 84% | TYLGY**S**YGTRI | VSTTH**D**PATPY | YDGTQ**H**TVVFQ |
| *M. chelonae* | 70% | NYLGY**S**YGTRI | VSTTN**D**PATPY | YEGTQ**H**TVVFQ |
| *M. abscessus* | 68% | TYLGY**S**YGTRI | VSTVH**D**PATPY | FDGTQ**H**TVVFQ |
| | | .********** | ***.:****** | : ********* |

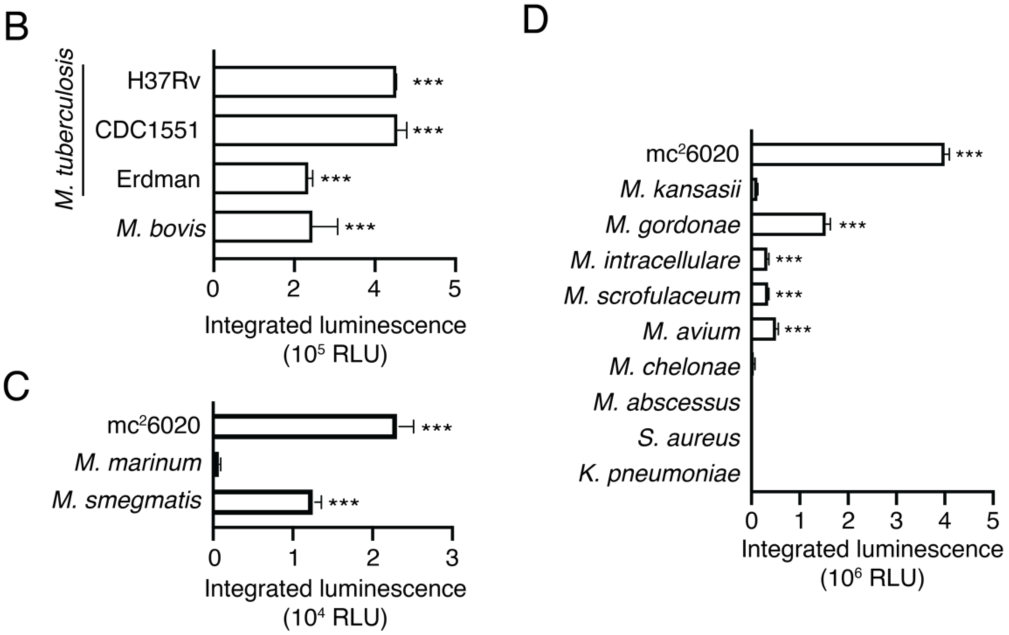


**Figure S4. Comparison of FLASH signal from NTMs and other bacteria.** (A) Sequence alignments for the three catalytic residues of Hip1 and putative homologs from NTMs. (B-D) Integrated luminescent signals for high density cultures (3x108 CFU/mL) of each bacterial species (mean ± s.d., n=3). H37Rv, CDC1551, Erdman, and mc$^2$6020 refer to *M. tuberculosis* strains. Each sample was compared to the no-bacteria control via one-way ANOVA with Dunnett's test (***, $p<0.001$).

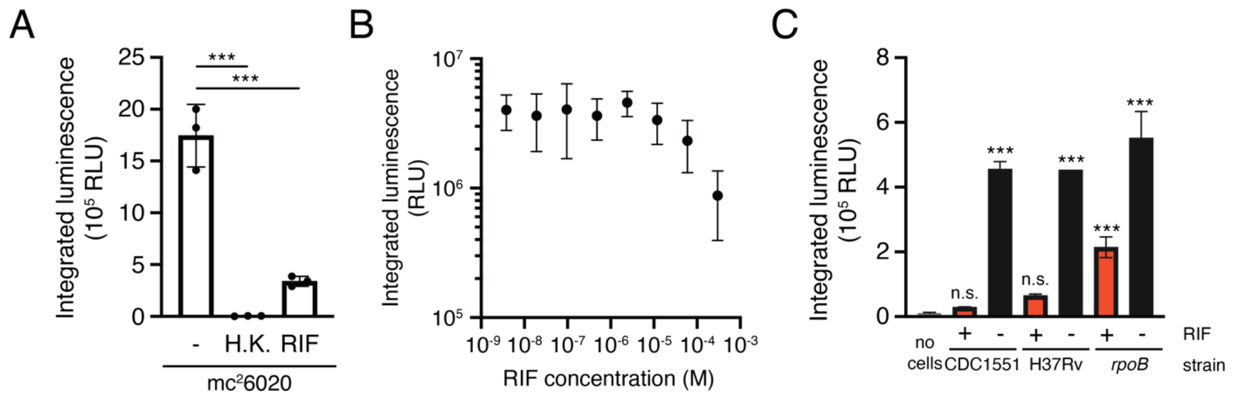


**Figure S5. FLASH and CTB measurements of RIF-treated cultures.** (A) FLASH signal resulting from untreated (-), heat-killed (H.K.), or 1.2 μM rifampicin-treated (RIF) *M. tuberculosis* mc$^2$6020 (mean ± s.d., n=3). Each sample was compared to live bacteria via one-way ANOVA with Dunnett's test (***, $p<0.001$). (B) Luminescence measurements of various concentrations of RIF in PBS immediately following treatment with AquaSpark. RIF decreases luminescent signal at high concentrations, but not at those relevant for *M. tuberculosis* killing. (C) Luminescent signal from CDC1551, H37Rv, or RpoB codon 526 mutant (*rpoB*) after 4 d of culture in the presence or absence of RIF. Samples are compared to the medium control via one-way ANOVA with Dunnett's test (***, $p<0.001$).

### Supporting Methods

#### Chemical Synthesis

All reactions requiring anhydrous conditions were performed under an argon atmosphere. All reactions were carried out at room temperature unless stated otherwise. Chemicals and solvents were either A.R. grade or purified by standard techniques. Thin layer chromatography (TLC): silica gel plates Merck 60 F254: compounds were visualized by irradiation with UV light. Column chromatography (FC): silica gel Merck 60 (particle size 0.040-0.063 mm), eluent given in parentheses. Reverse-phase high pressure liquid chromatography (RP-HPLC): C18 5u, 250x4.6mm, eluent given in parentheses. Preparative RP-HPLC: C18 5u, 250x21mm, eluent given in parentheses. Mass spectra were measured on Waters Xevo TQD. All reagents, including salts and solvents, were purchased from Sigma-Aldrich. Light irradiation for photochemical reactions: LED PAR38 lamp (19W, 3000K).

**Abbreviations. DCM** - Dichloromethane, **DMF** - N,N'-Dimethylformamide, **EtOAc** - Ethylacetate, **Hex** - Hexanes, **TFA** - Trifluoroacetic acid, **THF** – Tetrahydrofuran, **EEDQ** - *N*-Ethoxycarbonyl-2-ethoxy-1,2-dihydroquinoline, **TMSCl** - Trimethylsilyl chloride **HBTU -** Hexafluorophosphate benzotriazole tetramethyl uronium, **DIPEA** - *N,N*-Diisopropylethylamine , **DMBA - 1**,3-Dimethylbarbituric acid.

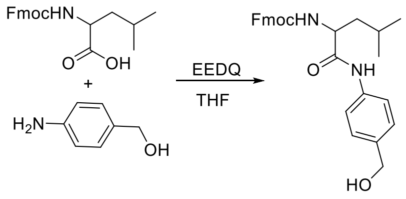


Compound **1** - L-Leu
Compound **2** - D-Leu

#### Compound 1 and 2

Compound **1** and compound **2** were synthesized according to a previously reported procedure *(53).*

#### General procedure 'A'

To a solution of the corresponding benzyl alcohol (247mg, 0.54 mmol) and NaI (246mg, 1.64 mmol) in ACN (15 mL) was added TMSCl (312µL, 1.64 mmol) at 0 °C. The mixture was brought to room temperature and stirred for 60 minutes and monitored by TLC (Hex:EtOAc 70:30). After full consumption of starting material, the reaction mixture diluted with EtOAc (100 ml) and was washed with brine (50 ml). The organic layer was separated, dried over $Na_2SO_4$ and evaporated under reduced pressure. Then, to the crude residue were added phenol enol ether *(54)* (200 mg, 0.48 mmol) and $K_2CO_3$ (138 mg, 1.0 mmol). The reaction was stirred at room temperature 30 minutes and monitored by TLC (Hex:EtOAc

70:30). After full conversion of the enol ether starting material, the reaction mixture was diluted with EtOAc (100 ml) and was washed with brine (50 ml). The organic layer was separated, dried over $Na_2SO_4$ and evaporated under reduced pressure the product was purified by column chromatography.

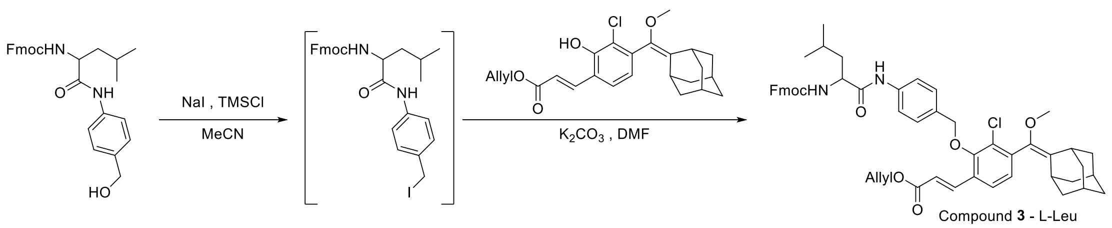


Compound **3** - L-Leu
Compound **4** - D-Leu

**Compound 3**

Compound **3** was prepared by general procedure ’A’ using compound **1** as starting material. Compound **3** was obtained in 67% yield as an off-white powder. MS (ES+): m/z calc. for $C_{52}H_{55}ClN_2O_7$: 854.4; found: 585.6 $[M+H]^+$.

**Compound 4**

Compound **4** was prepared by general procedure ’A’ using compound **2** as starting material. Compound **3** was obtained in 71% yield as an off-white powder. MS (ES+): m/z calc. for $C_{52}H_{55}ClN_2O_7$: 854.4; found: 585.5 $[M+H]^+$.

**General procedure ’B’**

The corresponding enol ether (compound **3** or **4**, 50 mg, 0.06 mmol) and piperidine (70 µL, 0.6 mmol) were dissolved in DMF (5 mL). The solution stirred for 30 minutes at room temperature and monitored by RP-HPLC (gradient of ACN in water). After full deprotection of the Fmoc was observed the solvent was removed under reduced pressure and the crude was dissolved in EtOAc was washed twice with 0.1M HCl (50 ml) and brine (50 ml). The organic layer was separated, dried over $Na_2SO_4$ and evaporated under reduced pressure. Then, the crude was added a premixed DMF (5 mL) solution containing tri peptide (42 mg, 0.07 mmol), HBTU (33 mg, 0.26 mmol) and DIPEA (22 µL, 0.36 mmol). The reaction was stirred for 60 minutes at room temperature and monitored by RP-HPLC (gradient of ACN in water). After completion, the reaction mixture diluted with EtOAc (100 ml) and was washed with 0.1M HCl (50 ml) and brine (50 ml). The organic layer was separated, dried over $Na_2SO_4$ and evaporated under reduced pressure. The crude product was dissolved in DCM (5mL) and was treated with DMBA (31 mg, 0.2 mmol) and $Pd(PPh_3)_4$ (11 mg, 0.01 mmol). The reaction was stirred at room temperature and monitored by RP-HPLC. Upon full deprotection of the allyl protecting groups the solvent was concentrated under reduced pressure and the product was purified by preparative RP-HPLC (gradient of ACN in water).

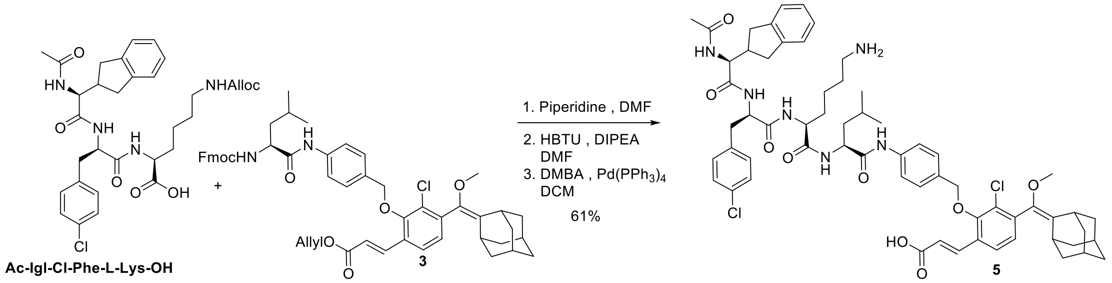

### Fmoc-Solid Phase Peptide Synthesis

Tripeptides **Ac-Igl-Cl-Phe-D-Lys-OH** and **Ac-Igl-4-Cl-Phe-D-Lys-OH** were synthesized using standard solid-phase peptide synthesis. Rink amide resin (282 mg, 0.17 mmol) was loaded into a filter cannula and swollen in DCM for 20 minutes. Resin was washed with DMF twice and deprotected in 20% (vol/vol) piperidine/DMF twice for 20 min and washed three times with DMF. Five eq of Fmoc-L-Lys(Boc)-OH (0.96 mmol, 289 mg) or Fmoc-D-Lys(Boc)-OH (0.96 mmol, 289 mg) were activated with 5 eq of HATU (0.96 mmol, 321 mg) and 5 eq of 2,4,6-collidine (0.96 mmol, 113 µL) in DMF for five min. The activated amino acid was added to the resin in DMF and allowed to react for 16 h, rotating at 25 °C. The resin was washed three times with DMF, two times with DCM, and the coupling reaction for the first amino acid was repeated for another 24 h. The resin was then washed three times with DMF, two times with MeOH and dried under vacuum. Before coupling a new amino acid, Fmoc-deprotection was performed two times in 20% piperidine/DMF for 30 min, washed two times with DMF, two times with DCM and taken up in DMF. Subsequent couplings for the P3 and P4 positions were accomplished using 5 eq of each amino acid together with 5 eq HBTU and 5 eq DIPEA in DMF for 16 h. Amino acids were Fmoc-Phe(3-Cl)-OH, and Fmoc-2-indanylglycine-OH. After each coupling the resin was washed two times with DMF, two times with DCM and taken up in DMF. After attachment of the P4 amino acid, the deprotected amino-terminus was capped using 5 eq acetic anhydride, 5 eq DIPEA in DMF. Finally, the peptide was cleaved from the resin and deprotected in 95% TFA, 2.5% TIS, 2.5% water and then precipitated in Et2O at -20°C. The precipitate was dissolved in 50% ACN/water and the product was purified by HPLC on a C18 column and lyophilized.

### Compound 5

Compound **5** was prepared by general procedure 'B' using compound **3** and **Ac-Igl-Cl-Phe-L-Lys-OH** (was synthesized via Fmoc-Solid Phase Peptide Synthesis) as starting materials. Compound **5** was obtained in 81% yield as a white powder. MS (ES+): m/z calc. for $C_{62}H_{74}Cl_2N_6O_9$: 1116.5; found: 1118.0 $[M+H]^+$.

HPLC analysis of Compound **5** (gradient ACN in water – 70%-100% ACN)

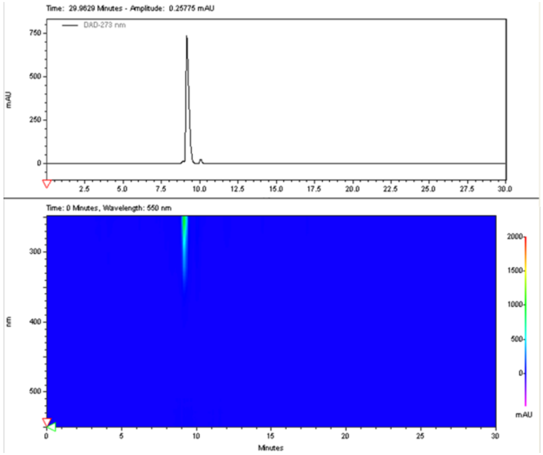


MS Spectra of Compound **5**

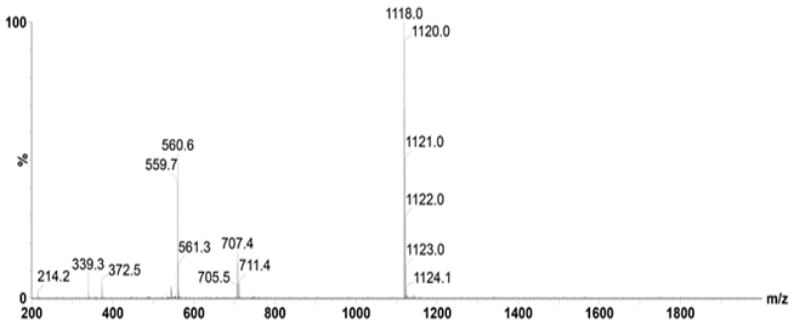


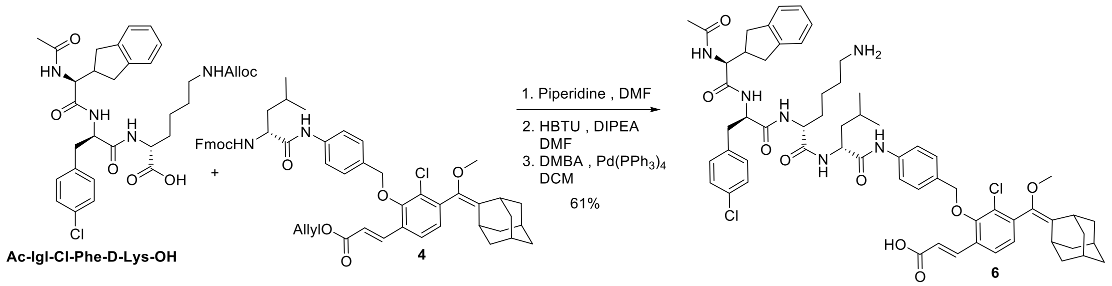


### Compound 6

Compound **6** was prepared by general procedure 'B' using compound **4** and **Ac-Igl-Cl-Phe-D-Lys-OH** as starting materials. Compound **6** was obtained in 64% yield as a white powder. MS (ES+): m/z calc. for $C_{62}H_{74}Cl_2N_6O_9$: 1116.5; found: 1118.0 $[M+H]^+$.

<u>HPLC analysis of Compound **6**</u> (gradient ACN in water – 30%-100% ACN)

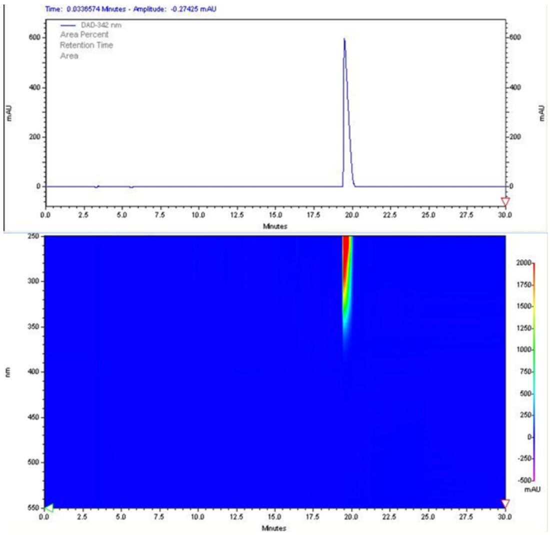

<u>MS Spectra of Compound **6**</u>

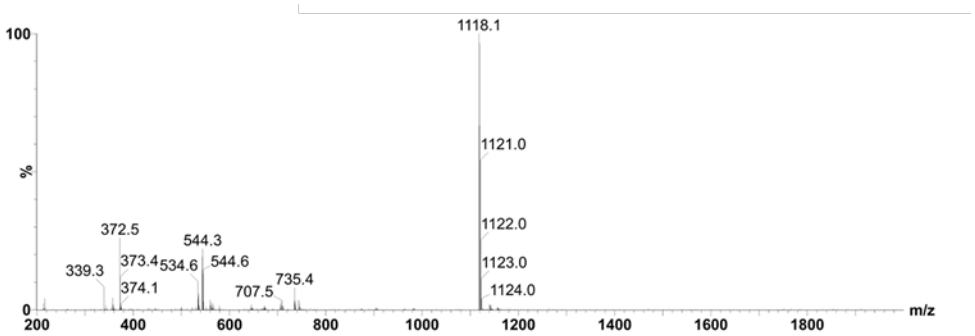


**General procedure 'C'**

The corresponding peptide protected enol ether (compound **5** or **6** - 20 mg, 0.02 mmol) was dissolved in DCM (20mL) and a catalytic amount of methylene blue was added to the mixture (~1 mg). Then, oxygen was bubbled through the solution while irradiating with yellow light. The reaction was monitored by RP-HPLC (gradient of ACN in water). Upon completion, 15 min, the solvent was concentrated under reduced pressure and the product was purified by preparative RP-HPLC (gradient of ACN in water).

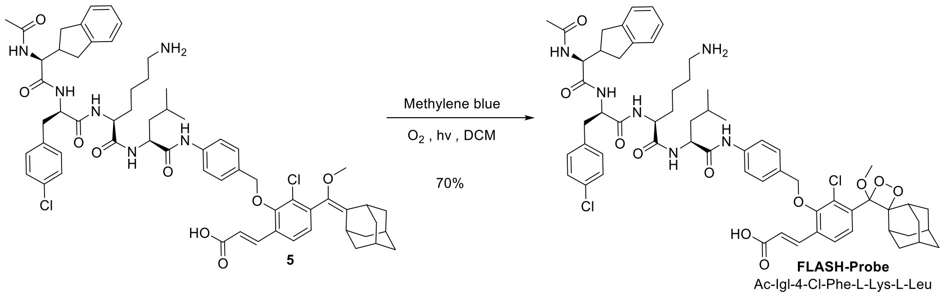


**FLASH-Probe (Ac-Igl-4-Cl-Phe-L-Lys-L-Leu)**

**FLASH-Probe** was prepared by general procedure 'C' using compound **5** and as starting material. **FLASH-Probe** was obtained in 70% yield as a white powder. MS (ES-): m/z calc. for $C_{62}H_{74}Cl_2N_6O_{11}$: 1148.5; found: 1148.1 $[M-H]^-$.

HPLC analysis of **FLASH-Probe** (gradient ACN in water – 50%-100% ACN)

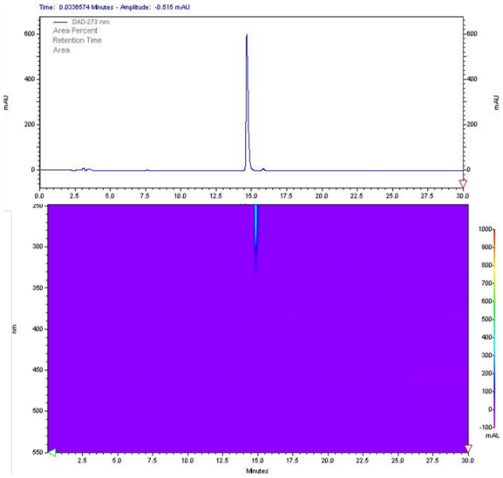

MS Spectra of **FLASH-Probe**

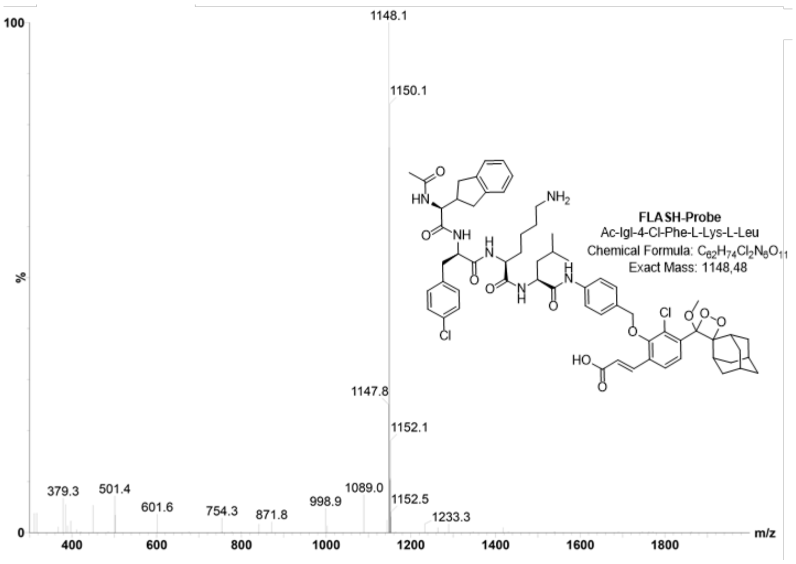


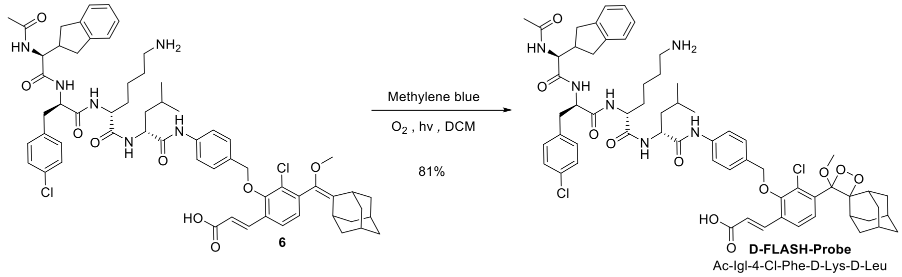


**D-FLASH-Probe (Ac-Igl-4-Cl-Phe-D-Lys-D-Leu)**

**D-FLASH-Probe** was prepared by general procedure 'C' using compound **6** and as starting material. D-**FLASH-Probe** was obtained in 81% yield as a white powder. MS (ES-): m/z calc. for $C_{62}H_{74}Cl_2N_6O_{11}$: 1148.5; found: 1148.1 $[M-H]^-$.

HPLC analysis of **D-FLASH-Probe** (gradient ACN in water – 50%-100% ACN)

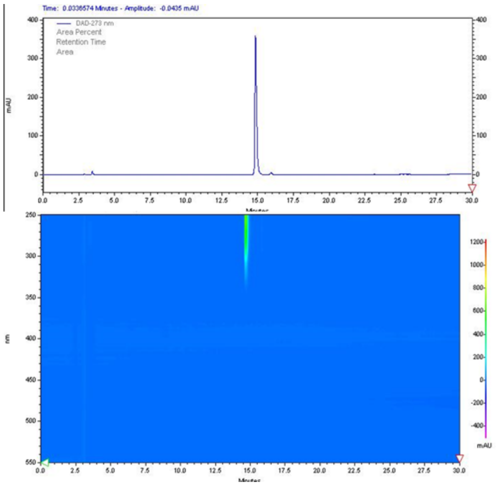


MS Spectra of **D-FLASH-Probe**

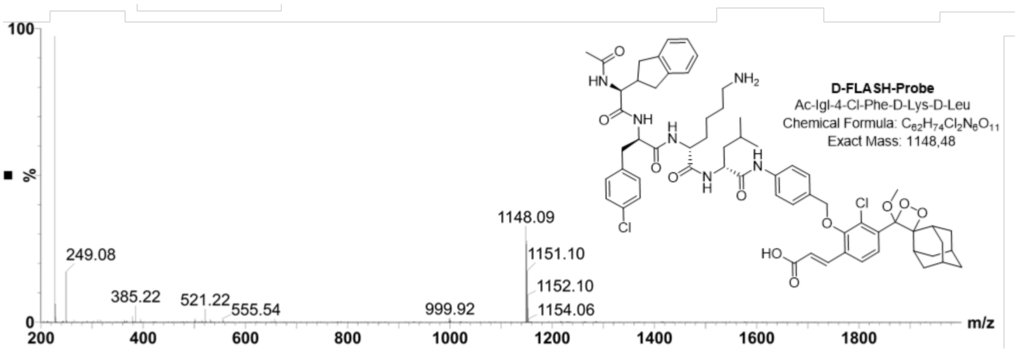
